# Isoswitching in human brain development and aging

**DOI:** 10.1101/2025.05.05.652255

**Authors:** Beril Erdogdu, Hyun Joo Ji, Zoe C. Rudnick, Mihaela Pertea, Steven L. Salzberg

## Abstract

Learning, reasoning, and working memory functions are attributed to the dorsolateral prefrontal cortex (DLPFC), a brain region that is highly evolved in primates and notably variable among individuals. Environmental and genetic factors likely contribute to this variability, but little is known about how they influence changes within an individual brain across the lifespan as different cognitive tasks and challenges arise. Most genetic studies focus on DNA mutations or changes in overall gene expression levels. However, genes can also alter the form in which they are expressed through alternative splicing. Using RNA-seq data from prenatal and postnatal human DLPFCs, we observed that many genes undergo a dramatic rewiring of their isoform usage around the time of birth. Further, thousands of genes continue to undergo gradual, temporally regulated shifts in their preferred isoforms, a phenomenon we term ‘isoswitching’. We present isoswitching as a major force in brain development and aging, capable of accurately predicting subject age from prenatal stages through adulthood and beyond eighty years of age. This study represents the first human brain age prediction based solely on RNA-seq data, establishing a molecular framework for understanding normative brain development and aging. We also report isoswitching in the brain of a closely related primate, the rhesus macaque.

## Introduction

From embryonic neurogenesis to senescence, the human brain evolves continuously through a precise orchestration of multiple cellular processes such as proliferation, axonal growth and pathfinding, selective cell death, and synaptic modulation(1, 2). Although these processes unfold in great complexity across both spatial and temporal dimensions, they are tightly regulated by genetic mechanisms that support the development and maintenance of a healthy brain(3, 4). These genetic mechanisms control processes as diverse as a child’s initial acquisition of language and the gradual decline of memory with advancing age, despite all cells within an individual retaining essentially the same DNA sequence throughout life. The regional and temporal specificity of brain function is largely achieved through variations in gene expression, involving both transcriptional and post-transcriptional regulatory mechanisms of RNA(5).

The generation and utilization of alternative isoforms (transcripts) from the same gene is one mechanism known to drive functional diversity in the human brain across different life stages(6). Differential transcript usage (DTU) refers to the change in the isoform preference for a given gene, without necessarily changing the total gene expression, and it plays a critical role in determining cell fate, function, and regulation across most eukaryotic tissues. The human brain exhibits a remarkably high rate of alternative splicing compared to other tissues, yet the corresponding DTU mechanisms and their functional consequences are still unknown(6-8). Two major developmental stages in the human brain are marked by substantial shifts in the transcriptional landscape: first, during prenatal development, the brain transitions from rapid cell division and differentiation to specialized, organ-specific gene expression; and second, during early postnatal life, brain maturation occurs in coordination with environmental inputs(4). Despite its clear relevance, the relationship between differential transcript usage and the processes underlying brain development and aging remains largely uncharacterized.

In our study, we first examine the developmental implications of this phenomenon by comparing prenatal and postnatal human brain samples. Integrating RNA sequencing data from 341 human dorsolateral prefrontal cortex (DLPFC) samples sequenced at the Lieber Institute for Brain Development (LIBD), we present the first large-scale comparison between the prenatal and postnatal human brain transcriptomes, revealing thousands of genes where the dominant isoform shifts entirely between prenatal developmental stages and post-birth. Among these is *SNAP25*, a gene encoding a protein essential for synaptic plasticity and neurotransmitter release, known to produce two major variants, *SNAP25a* and *SNAP25b(9)*. Previous studies in murine organisms have demonstrated that the developmental transition from *SNAP25a* to *SNAP25b* is pivotal for cognitive development and overall fitness for survival(10-14). Consistent with these findings, prior work shows *SNAP25b* to be the dominant isoform in adult human brain(15). Here, we extend these observations by providing a continuous temporal characterization of the *SNAP25* isoform transition in the human brain from prenatal to early postnatal stages.

The DLPFC is one of the most advanced regions of the human brain, responsible for problem solving, attention, and working memory. It is also the most recently evolved brain region, with an extended maturation period that continues well into early adulthood(16, 17). This prolonged developmental timeline distinguishes humans from other species, highlighting both intriguing similarities and important differences in neurodevelopmental processes. Examining the most significant of the genes implicated in DLPFC development, we found that isoform shifts have a clear correlation with brain aging throughout all stages of human life, extending well beyond prenatal and early postnatal development. Here, aging refers broadly to changes that occur across the lifespan and does not necessarily imply functional decline.

Using random forest regression models trained on the isoform usage matrix, we demonstrate for the first time that isoform usage patterns in the human brain consistently predict brain age (R² > 0.90) in multiple datasets, beginning from prenatal development and continuing through late adulthood, beyond 85 years of age. The simple and interpretable structure of the random forest models allowed us to identify over 5,000 genes as important contributors to brain age prediction. Further, we augmented the random forest predictor with lasso regressors specific to different age groups in a stacked model approach, enabling us to evaluate which genes are particularly important at each life stage. While we explore the functional relevance of a selected subset of these important genes, including *PALM*, *ARL16*, *RRAGB*, and *AGAP3*, the remaining genes are presented as valuable candidates for future investigation.

In support of our findings, we further validated the predictive power of isoform shifts using an entirely independent dataset from Cardoso-Moreira *et al*.(4) and extended this validation to a closely related primate, the rhesus macaque. Together, our results show that isoform shifts demonstrate widespread transcriptional modulation in the human brain throughout life and across the majority of genes. We refer to this dynamic reconfiguration of isoform dominance over time as ‘isoswitching’, and suggest that it represents a regulatory mechanism supporting healthy brain development, with potential implications for developmental dysregulation and brain aging.

## Results

The Lieber Institute for Brain Development houses an extensive collection of postmortem brain samples, spanning various life stages from prenatal development through infancy, adolescence, adulthood, and up to 85 years of age. We used RNA-seq data from this collection that was analyzed in earlier studies(18, 19) to explore the transcriptomic evolution of the human brain across different developmental stages and aging. Our primary analysis includes data from a total of 341 dorsolateral prefrontal cortex (DLPFC) samples, comprising 56 prenatal and 285 postnatal samples with no known history of psychiatric diseases. The dataset includes both male and female individuals, primarily of African American and Caucasian descent (**Figure 1a**).

**Figure 1.**
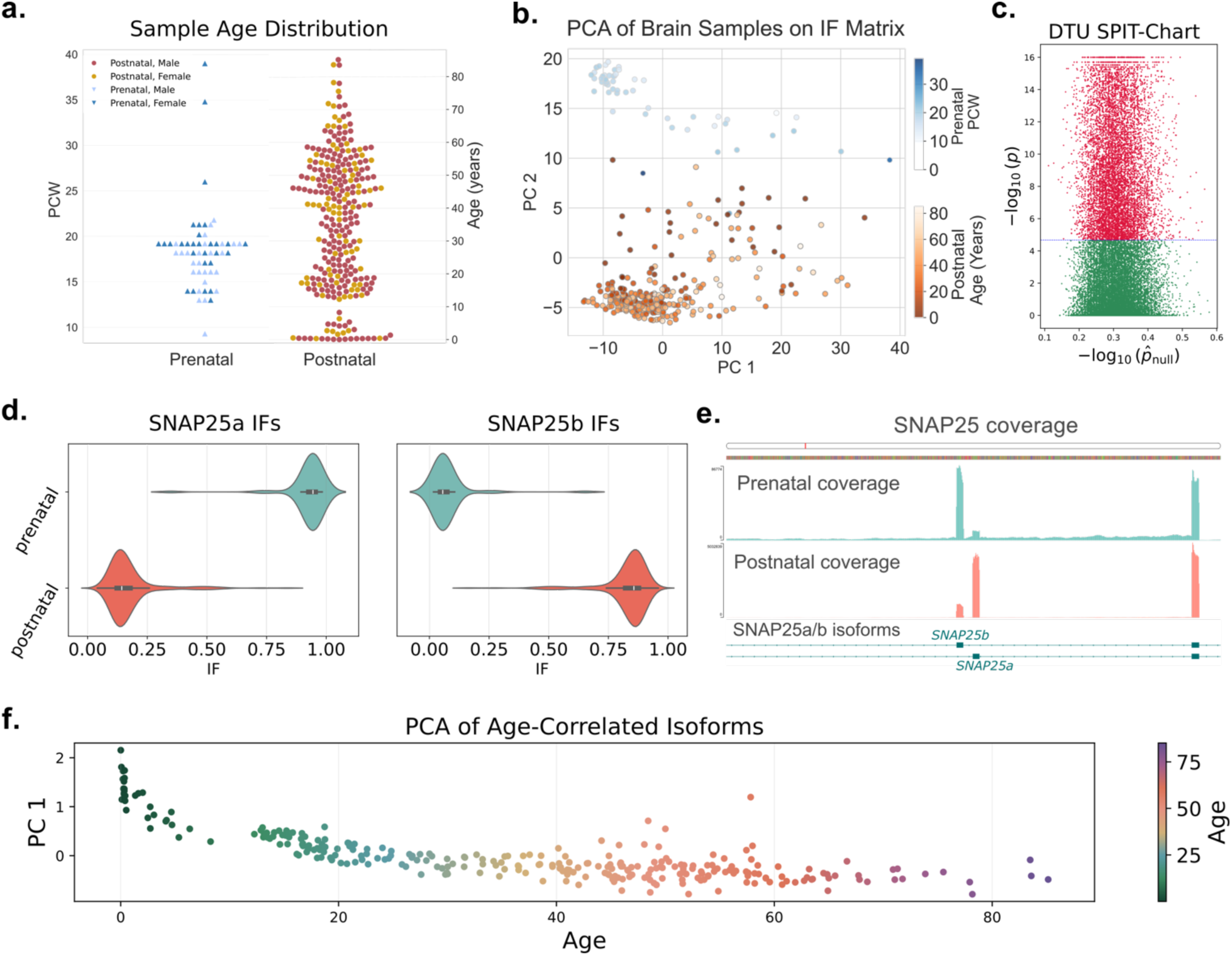
**a.** Age distribution of all LIBD samples, separated by developmental group and colored by sex. Postnatal sample ages are shown in years (right y-axis), while prenatal sample ages are shown in post-conception weeks (PCWs, left y-axis). **b.** Principal Component Analysis (PCA) of the isoform fraction (IF) matrix for all LIBD brain samples. PC1 and PC2 are shown. Prenatal samples are colored in shades of blue, with darker shades indicating proximity to birth. Postnatal samples are colored in shades of orange, with darker tones near birth and lighter tones representing increasing age. **c.** SPIT-Chart from the DTU analysis comparing prenatal and postnatal groups. Each transcript is represented as a point. The y-axis shows the –log10 (*p*-value) from the comparison, while the x-axis shows the null distribution from comparisons between random subsets of the control group. Significant transcripts are shown in red, and non-significant in green. **d.** Violin plots showing the distribution of isoform fraction (IF) values for the *SNAP25* variants across prenatal and postnatal samples. **e.** Coverage plot of SNAP25 switching dominant exons based on alignments of RNA-seq reads from prenatal (green, top) and postnatal (orange, bottom) samples. **f.** PCA of the IF matrix for DTU transcripts correlated with age. PC1 variation with age is displayed, with samples colored by age in years.

The role of alternative splicing in determining human cortical development has long been a subject of curiosity and investigation(20-23). Differential transcript usage provides a simplified snapshot of complex alternative splicing mechanisms, capturing the most prominent of changes in isoform expression. As our initial step of analysis, we looked into obvious patterns of isoform dominance within different developmental groups of the LIBD brain samples. Isoform fractions (IF) for transcripts in the CHESS 3 gene catalog(24) were computed for all samples using the quantification of RNA-Seq reads. The IF value represents the ratio of total gene expression that any isoform accounts for in a sample; e.g., if an isoform accounts for 80% of the expression of a particular gene, its IF will be 0.80. To help minimize noise, we refined our analysis to focus on a reliable set of genes and transcripts with sufficient expression evidence (see Isoform Quantification and Filtering section in Methods). A total of 29,757 transcripts from 9,645 genes were compared across samples after filtering. Upon basic dimensionality reduction with Principal Component Analysis (PCA), we observed a striking separation between the prenatal and postnatal groups (**Figure 1b**), suggesting systematic isoform shifts around the time of birth. Samples near the birth transition exhibited a weaker variance signal, positioning them closer to the opposite group; i.e., prenatal samples with higher post-conception weeks (PCW) tended to appear closer to early postnatal samples (infants near age 0). Analysis of variance revealed a significant association between developmental stage (pre- vs. postnatal) and PC2, with no significant associations observed for sex, race, or RIN (Supplementary Figure 1).

To further investigate the differential transcript usage (DTU) events between the two groups, we conducted DTU analysis on the prenatal and postnatal samples using SPIT(25). Developed for heterogeneous datasets and complex traits, SPIT is a conservative DTU detection tool that accounts for potential subgroups within cohorts. This ensures that changes affecting only a subset of a heterogeneous group, such as children or adolescents within the postnatal cohort, are not overlooked. We controlled for sex and ancestry as potential confounding factors, and multiple testing correction was performed within the SPIT framework using an empirical null distribution derived from randomized comparisons. After excluding genes influenced by these variables, SPIT reported 12,374 transcripts from 5,443 genes involved in significant DTU events between prenatal and postnatal brain samples, representing nearly half of all genes analyzed for DTU after filtering (**Figure 1c**). Among these, 311 genes were flagged as having highly significant associations with developmental stage.

*SNAP25*, a t-SNARE regulator of neurotransmitter release, displayed some of the most striking shifts between the two groups, showing nearly opposite IF distributions for its two isoforms, *SNAP25a* and *SNAP25b* (**Figure 1d**). The protein product of *SNAP25* is one of three proteins involved in the SNARE complex, which joins vesicles to the plasma membrane during exocytosis(26). In higher eukaryotes, two isoforms of *SNAP25* exist, *SNAP25a* and *SNAP25b*. The associated protein products of the two Isoforms are nearly identical, differing in only nine out of 206 amino acids(27). The isoforms have similar functions; however, *SNAP25b* has been shown to release a larger pool of vesicles, thereby increasing the potential of exocytosis in cells and leading to a more efficient facilitation of synaptic transmission(28).

The transition from the predominant expression of *SNAP25a* to *SNAP25b* from early adulthood onwards in mouse brain development is well-established and supports our findings(9-11, 15, 27, 29). In adult mouse brain, the developmentally correct expression of each isoform has been shown to be critical for synaptic maturation and neurotransmission plasticity. One study by Bark *et al*.(10) demonstrated that when targeted mutations are induced to impair the transition between isoforms, affected mice largely die between 3 and 5 weeks of age, the period during which *SNAP25b* expression becomes dominant. Later studies showed that mice lacking *SNAP25b* exhibit impaired learning and cognitive function in adolescence, but compensate for these deficits in adulthood(9, 14). While *SNAP25b* has been shown to be the primary isoform expressed in adult human brain(15), no studies have been done to indicate *SNAP25a* as the primarily expressed isoform in prenatal human brain or to investigate the temporal trajectory of the developmental isoform transition. In this work, we provide a comprehensive analysis of this transition.

Coverage patterns in the prenatal and postnatal samples, based on splice-aware RNA-Seq read alignment, highlight the precise exon switch that represents the transition from *SNAP25a* to *SNAP25b* (**Figure 1e**).

Along with *SNAP25*, we examined the remaining 310 DTU genes flagged by SPIT to assess whether sample age accounted for some of the heterogeneity observed in the postnatal group. We first selected transcripts whose IF values correlated with aging (indicated by a Spearman correlation threshold of |≥ 0.5|) and used their IF values to perform a new PCA. Separation of samples by age along PC1 is shown in **Figure 1f**.

### Isoswitching predicts brain age

In light of the sequential transition in isoform abundance patterns during development (**Figure 1b)** and the time-dependent variation observed in the postnatal samples (**Figure 1f**) based on the IF matrix of significant DTU genes, we asked whether these isoform shifts are temporally dynamic- exhibiting a continuous progression throughout life, including all prenatal and postnatal stages, rather than a discrete switch around the time of birth.

We began by analyzing the *SNAP25* transition in greater detail. The coverage plot in **Figure 2a** includes prenatal samples (S0) and further subdivides the postnatal group into developmental stages: ages 0–13 (S1), 13–25 (S2), 25–60 (S3), and 60+ (S4). Following the birth transition, *SNAP25a* expression gradually declines, while *SNAP25b* continues to rise, showing a clear trend in S0, S1 and S2 and a more subtle but persistent movement in S3 and S4. Also considering the temporal IF trajectories of variants *SNAP25a* and *SNAP25b* (**Figure 2b**), it is evident that their isoform dynamics are continuously shaped by time and show a clear association with sample age across the lifespan. Previous studies in model organisms have demonstrated this isoform shift; here, we describe its continuous dynamics and establish the timing of these changes in humans. We also examined *SNAP25* expression across tissues present in the Genotype-Tissue Expression (GTEx) V10 collection which offers RNA-Seq expression estimates from 19,788 samples across 54 tissues for most known human genes (30). *SNAP25* expression is largely confined to the brain, indicating that the observed isoform dynamics predominantly affect the nervous system (**Figure 2c**).

**Figure 2.**
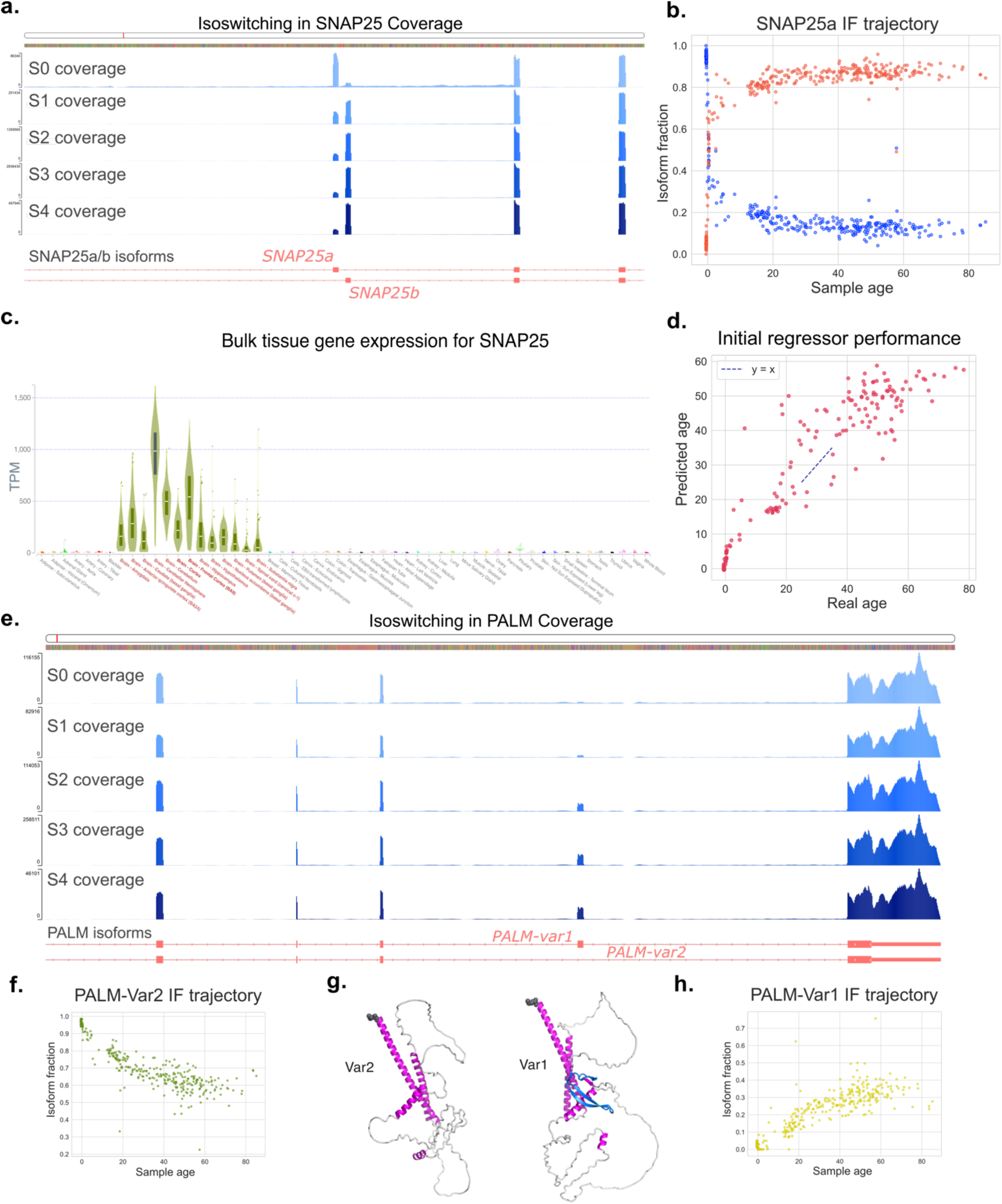
**a.** Coverage patterns of the shifting exons in *SNAP25*, based on alignments from all brain samples grouped as follows (top to bottom): ages 0–13 (S1), 13–25 (S2), 25–60 (S3), and 60+ (S4). Tracks are colored by age group, with darker shades representing older ages. **b.** Isoform fraction (IF) trajectories of the *SNAP25a* (blue) and *SNAP25b* (red) variants. Each sample is represented by two points, one per variant. **c.** Transcript per million (TPM) values for *SNAP25* across 54 human tissues from the GTEx project. **d.** Actual vs. predicted ages from the initial exploratory random forest model. **e.** Coverage patterns for exon 8 of the *PALM* gene for the same age groups in (a). **f.** Isoform fraction (IF) trajectory of *PALM* variant 2, which is dominant in prenatal samples. **g.** Protein structures of PALM variants. The additional beta sheets introduced by the merged exon are shown in blue. **h.** IF trajectory of *PALM* variant 1, increasingly favored in older samples, in the same layout as (f).

These findings led us to explore whether brain age could be predicted solely from the isoform fractions of a selected set of genes. Using the original set of 29,757 transcripts from 9,645 genes prior to any feature selection, we achieved a highly accurate prediction of brain age (Testing: R² = 0.85, RMSE = 8.62; Training: R² = 0.98, RMSE = 3.58) in our initial investigation (**Figure 2d**). Given the disproportionately large feature space relative to the number of samples, we opted for a simple learning framework and employed a random forest regressor(31-33) with 100 trees to avoid over-fitting. This exploratory model was trained on a randomly selected half of the LIBD brain samples, and tested on the other half. In this step, no more than half of the feature space was considered for each node split, and trees were grown to their maximum depth, producing homogeneous leaves to evaluate learning capacity (Average depth: 11.36, Min depth: 9, Max depth: 16). These parameters were later adjusted in the final model to facilitate interpolation and support feature selection.

Our primary objective in predicting brain age was to identify genes that undergo isoswitching with stable and well-defined trajectories across development and aging. These genes serve as valuable candidates for markers of brain aging, and the biological significance of their isoswitches provides insight into the mechanisms that regulate changes in the brain over time.

We assessed feature significance within the decision forests using the Gini score, which quantifies the total reduction in impurity contributed by each feature across all tree splits, normalized over the forest and the feature space(32, 33). Using this metric, Paralemmin-1 (*PALM*) emerged as the top-ranked gene, contributing the most significant splits in the model. Coverage patterns of *PALM* (**Figure 2e**) reveal a gradual emergence of an exon with minimal expression in the prenatal stage, becoming more pronounced in S1 and steadily increasing through later life stages (S2–S4). Examining the IF trajectories of the two prominent *PALM* variants reveals a nearly linear shift with age.

*PALM* encodes a phosphoprotein located on the cytoplasmic surface of plasma membranes that plays a role in the regulation of membrane dynamics. Previous studies have shown that alternative splicing of *PALM* regulates neurodevelopmental processes such as the maturation of dendritic filopodia into spines, and the trafficking and membrane localization of AMPA-type glutamate receptors (e.g., GluR1) and D3 dopamine receptors(34-36). It has been demonstrated that alternative splicing involving the inclusion of exon 8 creates enhanced effects in these processes, with deletional mutational analysis indicating that amino acids 154–230 of paralemmin, corresponding to exon 8, show the strongest interactions with full-length D3(36).

Consistent with previous findings, we observed isoswitching patterns supporting the alternative splicing of exon 8, defined with respect to the MANE(37) isoform of *PALM*, where the predominantly expressed isoform transitions as a function of age from *PALM* transcript variant 2 (CHESS 3 ID: CHS.25148.2, RefSeq(38) ID: NM_001040134.2) to *PALM* transcript variant 1 (CHESS 3 ID: CHS.25148.1, RefSeq ID: NM_002579.3) (**Figure 2f-h**). We also examined the structural differences between the two isoforms at the protein level using ColabFold(39), and found that exclusion of exon 8 results in the loss of four adjacent beta sheets, as shown in the protein structure of the prenatal-dominant isoform CHS.25148.2 (**Figure 2g**). Given that this region of the PALM protein exhibits the strongest inhibitory interaction with the D3 dopamine receptor, it is plausible that increased inclusion of this domain with age may contribute to a reduction in dopamine signaling. We present evidence for the full trajectory of this isoswitching across all stages of development in the human brain.

### Extending to a broad genetic landscape with predictive power

To extend our observations beyond individual examples like *PALM* and *SNAP25*, we next investigated how many genes with similarly strong and temporally consistent isoform shifts could be collected using our random forest regressor. Transcripts from these two genes were the top most significant features in the random forest model, respectively. After removing all isoforms of *PALM* and *SNAP25* and retraining the regressor on the same randomly selected half of the LIBD brain samples, the model maintained comparable predictive performance (R² = 0.83, RMSE = 9.31), with the top-ranking feature now corresponding to an isoform of *RECQL4*.

*RECQL4* is a member of the RecQ helicase family and belongs to an enzyme group that is crucial for DNA stability and repair. While the role of its isoforms in the nervous system is not well-studied, mutations in *RECQL4* have been directly associated with features of premature aging and an increased risk of cancer(40-42), and could also play a role in the alternative splicing of the its RNA molecule(43). **Figure 3a** shows a gradual increase with age in an isoform that includes a continuous C-terminal exon spanning regions that are intronic in the earlier-expressed isoform. We then removed *RECQL4* and applied feature ablation by recursively eliminating all isoforms of the most important gene in each iteration, continuing this process until the model reached an unacceptably low level of performance(44). Surprisingly, even after removing 5,000 genes, the model continued to perform well above random, retaining a meaningful level of predictive accuracy (**Figure 4a**). These findings indicate that isoswitching relevant to brain aging is not confined to a few key genes, but is broadly distributed across the transcriptome, with many genes carrying predictive value.

**Figure 3.**
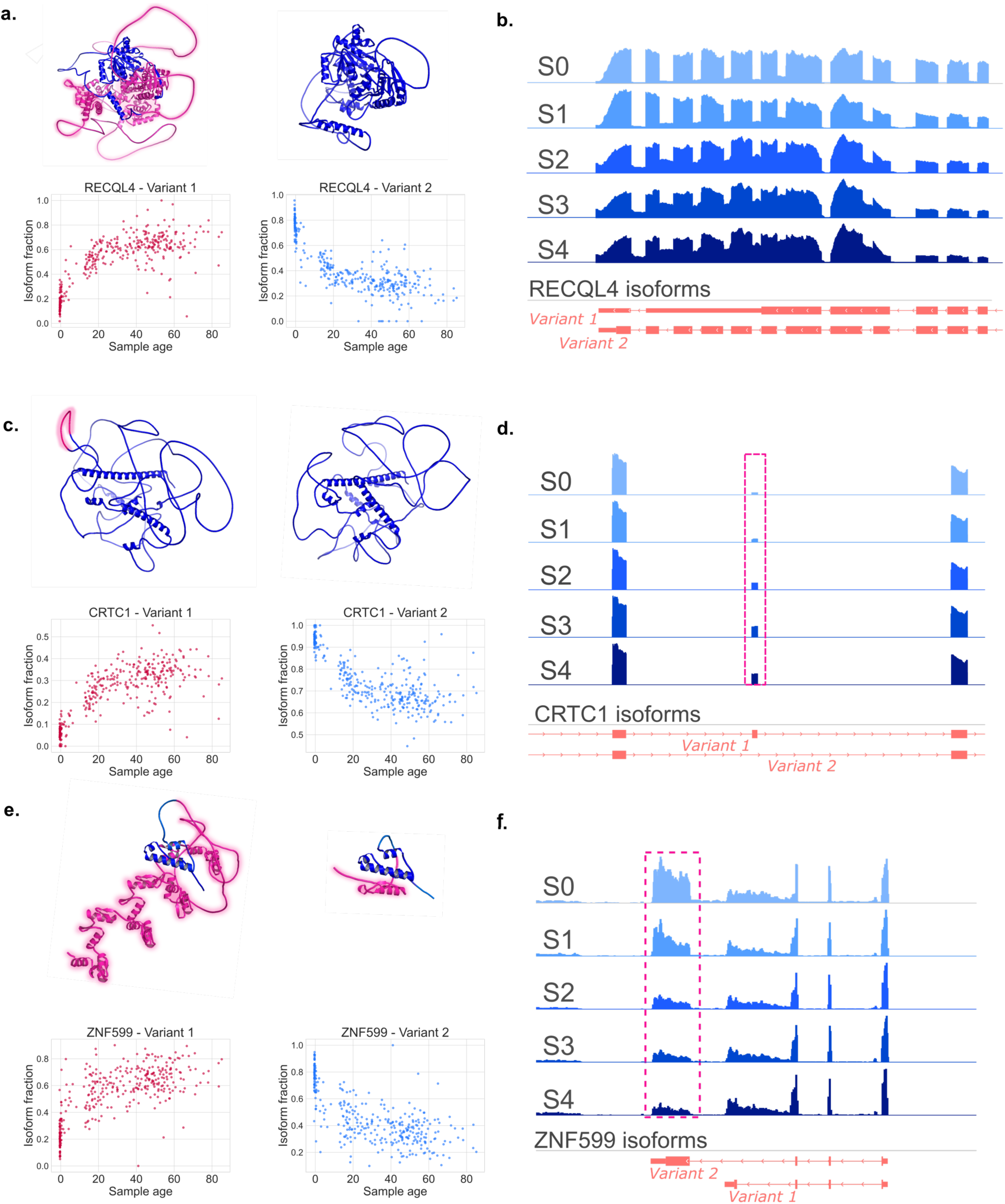
Examples of isoswitching genes that have not been previously characterized. Protein structure differences between the switching isoforms are shown (highlighted in pink) along with their temporal isoform trajectories. CHESS 3 IDs are provided for each variant. **a.** Protein products of switching isoforms of *RECQL4* (top row) with their corresponding isoform trajectories (bottom row). (Variant 1: CHS.54794.31, Variant 2: CHS.54794.1). **b.** Coverage patterns for the switching domain in *RECQL4*, as in Figure 2a,e. **c,d.** Same as (a,b) for *CRTC1* (Variant 1: CHS.26110.1, Variant 2: CHS.26110.2). **e,f.** Same as (a,b) for *ZNF599*. (Variant 1: CHS.26430.5, Variant 2: CHS.26430.1).

**Figure 4.**
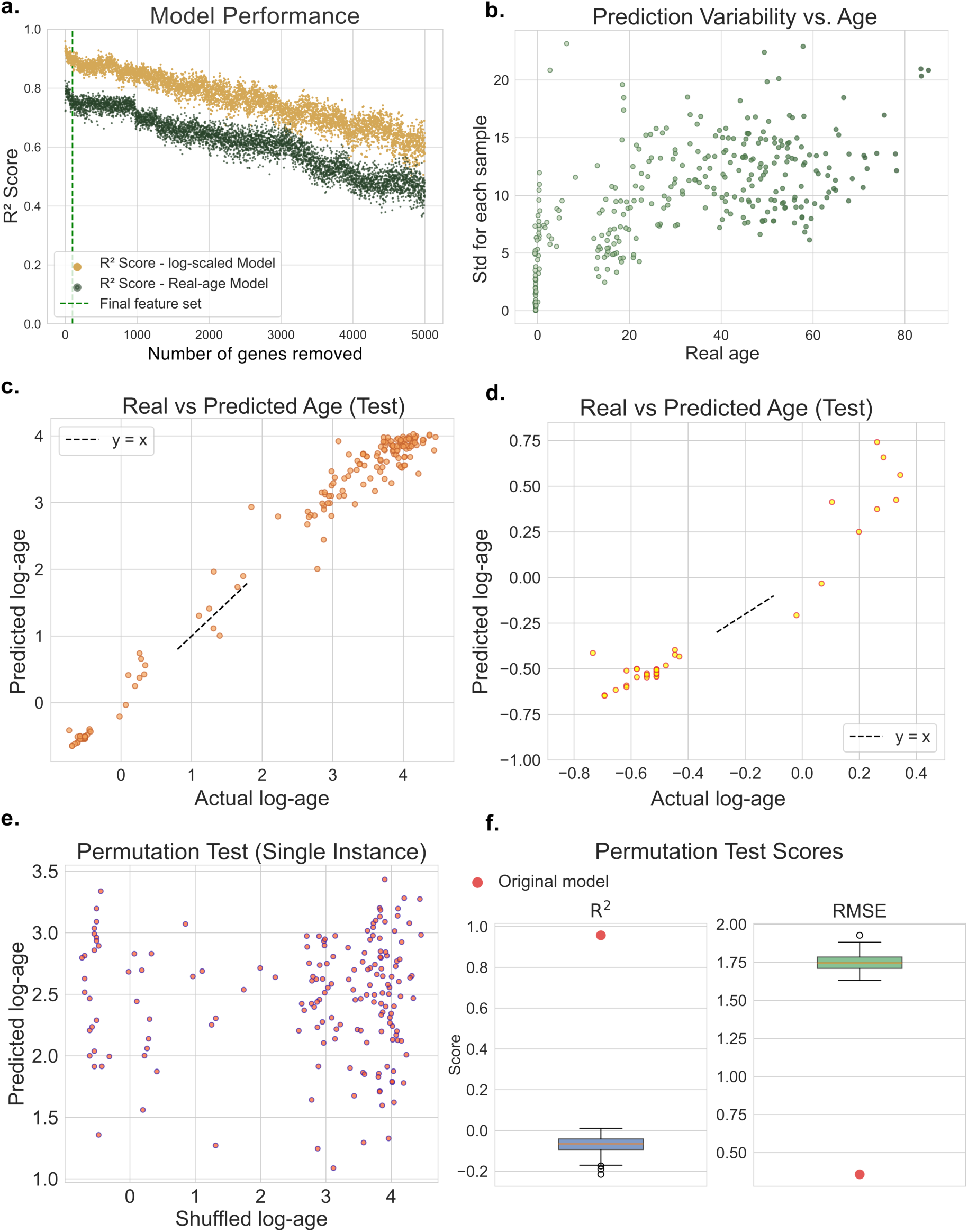
**a.** R^2^ scores throughout the feature ablation process. Log-transformed age (yellow) and raw age (green) models are shown. **b.** Standard deviation of predictions across the 100 trees in the random forest for each sample, plotted against sample age. **c.** Model performance for the random forest trained on log-scaled ages (predicted vs. actual log-age). **d.** Same as (c), zoomed to highlight prenatal and infant samples. **e.** Predictions from a random forest regressor trained with shuffled log-age values (single permutation instance). **f.** Box plots of R^2^ and RMSE for models trained as in (e) across 100 permutations. Corresponding scores from the original (non-shuffled) model are shown in red.

Unlike *PALM* and *SNAP25*, isoform shifting in most of these genes has not been previously characterized, and functional analyses at the transcript level are largely lacking. To highlight such previously unreported isoform dynamics, **Figure 3** illustrates the temporal trajectories of switching isoforms in *RECQL4*, *CRTC1*, and *ZNF599*, together with the protein structures that differ between their respective isoforms as predicted by ColabFold(39). CRTC genes have been studied for their roles in aging-associated cognitive decline at the level of gene expression; however, isoform-level regulation has not been explored. Our results bring this additional dimension into focus across development (45, 46). Literature on the role of *ZNF599* in brain function is limited, while the broader class of zinc finger proteins has been implicated in brain development and neurodevelopmental disorders(47). We provide a ranked set of the top 5,000 predictive genes as a resource of strong candidate genes for future mechanistic studies similar to those conducted on *PALM* and *SNAP25* (Supplementary Table 1).

### Scaling of brain age and time

A random forest regressor is an ensemble learning method composed of multiple decision trees, with 100 trees used in our models. Each tree independently generates a prediction for a given sample, and the final model output is calculated by averaging these individual predictions. To assess the model’s confidence and stability, we examined the variance among the predictions made by different trees for each sample, providing a measure of uncertainty in the decision-making process. The standard deviation of tree predictions for the LIBD brain samples (**Figure 4b**) shows a consistent increase in prediction uncertainty with age. This trend likely reflects two fundamental biological principles.

First, the biological significance of time varies substantially across different stages of life. A single year during prenatal development encompasses a dense sequence of cellular processes, including proliferation, migration, and differentiation. In contrast, a year in late adulthood is marked by slower biological change(48). The difference between a newborn and a one-year-old is profound, while the difference between a seventy-year-old and a seventy-one-year-old is comparatively subtle and less predictable. Second, while early development tends to follow a structured and genetically guided trajectory, aging is a highly individual process shaped by the interaction of genetic, environmental, and lifestyle factors. Unlike development, where milestones occur in a relatively uniform pattern, aging progresses at different rates in different individuals(49). Two people of the same chronological age may differ significantly in brain health, cognitive performance, and overall physical condition, reflecting the complexity and variability of aging(50, 51).

While we cannot fully account for individual differences in the progression of aging due to limitations in available data and scientific knowledge, we can adjust for the biological significance of time across different age groups. Given our objective of identifying meaningful gene candidates that undergo significant and well-defined isoform shifts predictive of age, it is crucial to recognize that the biological impact of the same time unit is not uniform across the lifespan. An error of one year should not be weighted equally for a 60-year-old individual and an infant, as this would undervalue genes that are crucial for prenatal and early brain development.

An intuitive transformation to address this issue is to predict age on a logarithmic scale rather than using raw age values. This approach assigns greater importance to age differences in early life while gradually reducing their impact in later stages. To ensure all values remain positive, ages were incremented by 1 before applying the natural logarithmic function. The random forest model trained on log-transformed ages (referred to as log-ages hereafter) appropriately assigns greater weight to errors made during earlier stages of life and achieves an exceptional R² score of 0.97 (log-age RMSE = 0.29). Training R² and log-age RMSE were 0.99 and 0.15, respectively. After feature selection (see Methods), using only the isoform fraction matrix from the top 100 selected genes, the model achieves an even higher R² score of 0.98 (**Figure 4c**, log-age RMSE = 0.27). For the feature-selected model the training R² and log-age RMSE were 0.99 and 0.13, respectively. Average tree depth was 12.21, with minimum and maximum depths of 10 and 15, respectively. Focusing on the prenatal and infant sample group (ages -1 to 1), we observe that isoswitching in these top genes not only predicts aging with high accuracy in postnatal stages, but also effectively tracks developmental progression in prenatal and infant samples based on post-conception weeks (**Figure 4d**).

### Assessing the Influence of Potential Confounders

When working with stratified biological data consisting of individuals from different backgrounds, there is always a risk that confounding variables influence the analysis. While available covariates were accounted for in the DTU analysis between pre- and post-natal samples, the random forest regressor does not incorporate this metadata. In addition, not all potential confounders are captured in the available sample annotations.

To assess the influence of strong confounders on model predictions, we performed a permutation test by randomly shuffling the age values of the samples and training the model on these permuted labels. Specifically, we retrained the model using the full feature matrix of IF values (29,757 transcripts) and randomly shuffled log-scaled ages for the LIBD samples. As shown in **Figure 4e**, the model trained on permuted ages exhibits very poor performance (R² = -0.09), suggesting that it does not capture strong signal from potential confounders.

We repeated this procedure across 100 random splits of the data and observed consistently poor performance. **Figure 4f** compares the R² and RMSE values from the original model (trained on the 29,757 transcripts and non-shuffled log-ages) to those obtained from the permutation test iterations. While we cannot fully rule out the presence of confounding factors, these results suggest that they are not a major driver of model performance in this setting.

Another consideration is that the observed changes in isoform abundances may reflect underlying changes in gene expression, which has been linked to aging in both model organisms and humans (52-56). To assess this, we built a separate model using total gene expression values instead of isoform abundances (Supplementary Methods, Supplementary Figure 3a,b). We then examined whether genes that change in expression across development and aging are the same genes that show predictive power through isoswitching events.

Upon applying feature ablation to the total gene expression model, we observed minimal overlap between the most predictive genes identified from isoform abundance and total expression. No genes were shared among the top 100 features, only five genes overlapped within the top 200 (*SEPTIN4, HAPLN2*, *CLDND1*, *SEPTIN8*, *GNG4*), and 65 genes were shared among the top 1000 (Supplementary Figure 3c,d, Supplementary Table 3). These results indicate that genes whose isoform usage is most predictive of age are largely distinct from those whose total expression levels carry age-associated signal.

While changes in total gene expression have been linked to aging, our results show that isoswitching provides an additional and largely non-overlapping source of predictive information. This suggests that isoform-level variation captures aspects of aging biology that are not fully reflected in gene expression alone, although these two factors may be biologically linked through shared regulatory mechanisms which remains to be determined.

### Developmental windows of gene relevance

Using our predictive model for estimating brain age across prenatal development, postnatal maturation, and aging, we identified a robust set of 100 genes that consistently contribute to its predictive accuracy. However, it is unlikely that all 100 genes contribute equally throughout the entire course of brain development and aging. Instead, distinct subsets are likely to be functionally relevant at different periods. For instance, a gene essential for neurogenesis and early cortical formation is unlikely to play a central role in the molecular processes associated with later-life decline, though some continuity between developmental and aging mechanisms may exist.

To better characterize the temporal dynamics of gene importance, we added an additional predictive layer on top of the random forest regressor. In this stacked modeling framework, the random forest first estimates the age of each sample. The predicted age is then used to assign the sample to one of five lasso regression models, each designed for a specific range of ages, designed to be overlapping: [-1, 1], [0, 10], [1, 25], [10, 60], and [25, 100].

This framework serves two complementary purposes. First, it allows us to segment human brain development and aging into biologically meaningful time periods that may correspond to distinct molecular transitions. Second, by using overlapping age ranges, the model provides a bridge between adjacent life stages, capturing smooth transitions rather than treating developmental phases as discrete, isolated units. Unlike prenatal samples -whose isoform profiles are strikingly distinct from postnatal ones- and the most elderly individuals, many samples fall within the overlapping range of two lasso regressors, allowing for interpolation or extrapolation from both models.

For instance, a sample with a predicted age of 20 years from the random forest is processed by both the [1, 25] and [10, 60] lasso models, receiving predictions from each (for details see Methods, Stacked model). This overlap enables direct comparison between models trained on different temporal windows, allowing for a comparison of predictions depending on the chosen reference point in the temporal axis.

To identify the most influential genes, we conducted an exhaustive pairwise evaluation. From the set of 100 top-ranked genes, we considered all 4,950 possible gene pairs {*g*_*u*_, *g*_*v*_} where *u* ≠ *v*. For each gene pair, we trained and tested the lasso layer using only the isoform fraction vectors corresponding to transcripts from those two genes. Regularization parameter *λ* was set to 0.05 to account for the reduced feature space. The resulting Spearman correlation score was compared to that of the original model trained on all 100 genes. Gene pairs that produced minimal loss in correlation were interpreted as being highly informative on their own, suggesting they possess strong and independent predictive value for the corresponding range of brain age. Using this method, we identified genes that, when included in a randomly selected pair, preserved model accuracy nearly as well as the full 100-gene model. These genes represent the most robust contributors to age prediction across distinct developmental periods, appearing as prominent red tracks in the heatmaps shown in **Figure 5a**. As the age categories widen and shift toward older groups across the lasso models from *l*_0_ to *l*_4_, prediction ambiguity increases, resulting in a gradual decline in the number of such stand-alone predictor genes.

**Figure 5.**
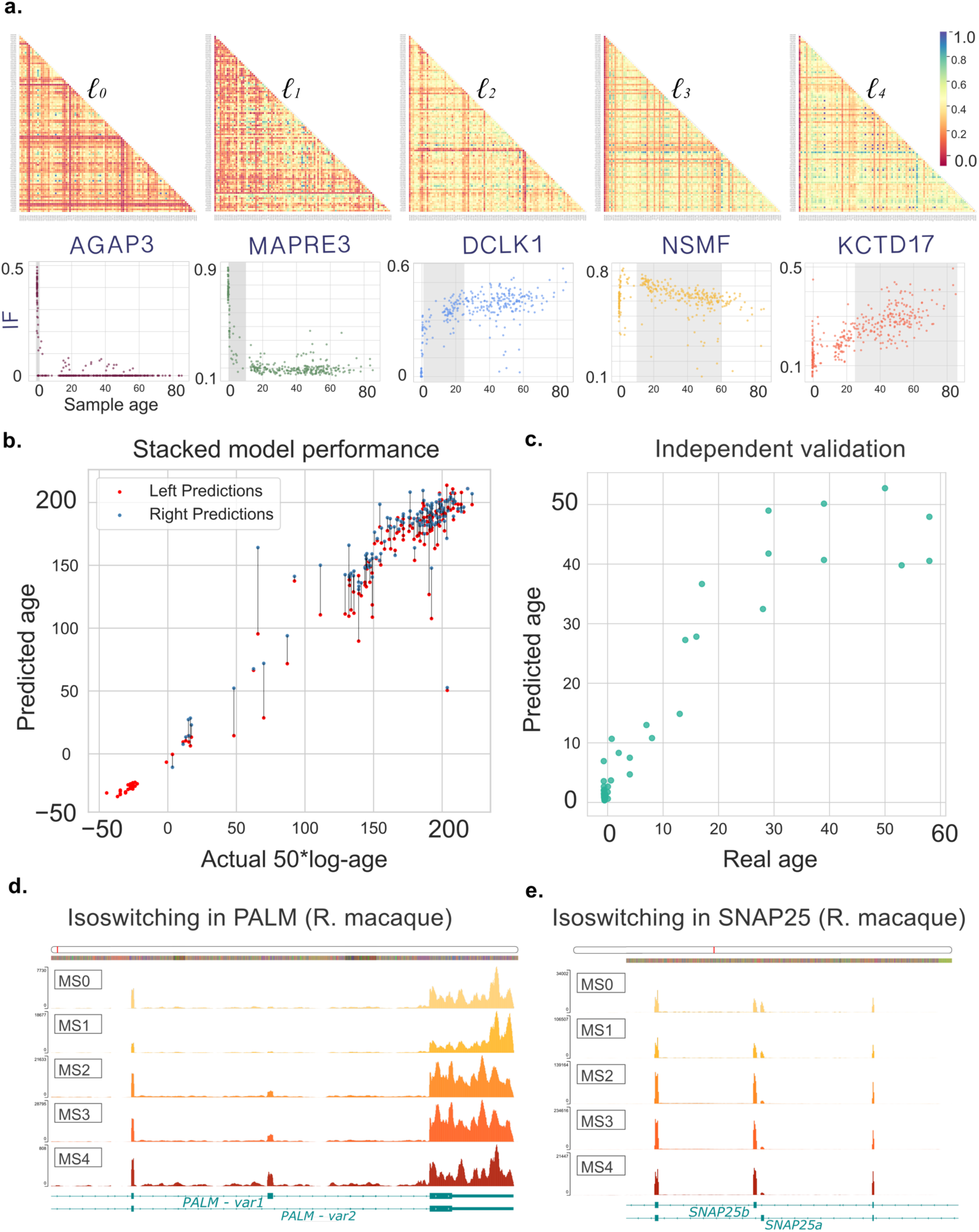
**a.** Heatmap of drops in accuracy for lasso regressors when trained on every possible pair of genes from the top 100. Lower drops (red) indicate stronger predictive power. For each lasso model, one gene from the red heatmap track was selected to illustrate an isoswitching variant trajectory, shown below each panel. The age range for the model is highlighted in gray. **b.** Performance plot for the stacked model trained on log-ages. For samples predicted by two lasso models, the prediction from the “younger” model is shown as the left (red) prediction and the “older model” as the right (blue), connected by a line. **c.** Performance of the random forest regressor on an independent dataset from Cardoso-Moreira et al.(4). **d.** Coverage patterns for exon 8 in *PALM*, present in *PALM* var1 but not var2, based on RNA-seq alignments from 32 rhesus macaque brain samples. Samples are grouped as follows: MS0 (prenatal), MS1 (ages 0–4), MS2 (ages 4–8), MS3 (ages 8–20), and MS4 (ages 20+). Tracks are colored by group, darkening with age. Introns are shortened for clarity. **e.** Same as (d), showing the isoform-switching region in *SNAP25*.

From the red tracks in the heatmaps, we manually selected a subset of stand-alone predictor genes and inspected their IF trajectories across all LIBD samples. One representative example from each heatmap is shown below its corresponding panel in **Figure 5a**. We observed a clear distinction in isoswitching behavior across developmental periods. In the lasso regressor trained on samples from the youngest subjects (prenatal to age 1), the most predictive genes exhibited a sharp, binary-like shift occurring precisely around the time of birth. In contrast, genes that contributed most to age prediction in later stages-such as those in the [10, 60] age range-showed gradual and continuous changes in isoform fractions over time. This difference suggests that some genes undergo abrupt regulatory transitions at birth, while others undergo gradual but sustained isoswitching that unfolds throughout life.

To evaluate the effectiveness of this framework, we applied it to the test set, where each sample was first processed by the random forest regressor and then routed to the appropriate lasso model for final age prediction. For samples that fell into overlapping age intervals and received predictions from two lasso models, the final estimate was calculated as the average of the two.

As expected, for such samples the lasso model trained on the younger age range almost always produced lower age estimates than the model trained on the older range (**Figure 5b**).

Importantly, no unexpected discrepancies were observed between adjacent lasso models, indicating strong consistency in predictions across overlapping age windows. The overall coefficient of determination (R²), based on the averaged predictions for all test samples, was 0.948 with scaled-up log-ages and 0.872 when using raw age values.

### Testing with independent data

Having established a model with strong predictive performance across the human lifespan, we next examined whether the observed relationship between isoform shifts and brain aging is robust and reproducible. To answer this question, we selected an independently published dataset of postmortem brain samples spanning a broad age range. We aimed to evaluate whether the model could accurately predict ages in an external cohort, and to determine whether the 100 key genes identified in our original analysis exhibited similar isoform fraction trends in a different population.

For this external validation, we selected the dataset published by Cardoso-Moreira *et al*.(4), which provided an ideal test case for assessing the robustness of our findings in an independent cohort. Cardoso-Moreira *et al*. conducted a study on organ development across seven species using bulk RNA-Seq data collected at multiple time points, spanning both prenatal and postnatal life stages. Their study examined gene expression changes in seven organs (forebrain, hindbrain, heart, kidney, liver, ovary, and testis) and explored the relationship between evolution and development. They found that gene expression patterns in organs tend to be more conserved across species during early development, with increasing divergence at later stages of life.

While the dataset includes only 53 human forebrain samples, with 32 being prenatal, it provides a valuable resource for studying developmental trajectories. Postmortem human brain data spanning a wide range of ages is exceptionally scarce, making this dataset particularly useful for validating our findings. We applied our random forest regressor-trained on the LIBD dataset and restricted to the top 100 predictive genes-to this external set of 53 samples, following the same quantification, normalization, and filtering procedures used for the LIBD RNA-Seq data.

Predicting log-scaled ages using the selected 100 genes yielded an R^2^ score of 0.862 (RMSE = 8.46). Similarly, applying the random forest regressor trained on the LIBD samples using real ages and the selected set of 100 genes resulted in strong performance, with an R² score of 0.84 (**Figure 5c**, RMSE = 6.59), compared to the R² of 0.86 (RMSE = 8.97) on the original LIBD test set. The stacked model predicted the log-ages (scaled up by a factor of 50, for details see Methods, Stacked Model) of the samples with an R^2^ of 0.92, comparable to its performance on the LIBD samples (R^2^ = 0.95).

We observed that the correlation between predicted and actual ages was noticeably lower for prenatal samples in this dataset. We attribute this to differences in the developmental timeframes represented in the two datasets. The LIBD dataset, on which our model was trained, captures a narrow window within prenatal development, whereas the Cardoso-Moreira *et al.*(4) dataset spans both earlier and later stages of pregnancy. As a result, our model had less data on the range of isoform fraction patterns and their changes in the earliest stages of development.

### Validation in the rhesus macaque brain

We next investigated whether similar patterns of isoswitching as a function of age could be observed in one of our closest relatives, the rhesus macaque. To explore this, we utilized the forebrain samples from the Cardoso-Moreira *et al*.(4) study, which included 32 macaque brain samples, nine of which were prenatal. The ages of these samples ranged from 93 post-conception days to over 20 years.

Unlike the well-annotated human transcriptome, the macaque transcriptome is considerably less well-characterized, with fewer documented alternative isoforms per gene. As a result, direct quantification using Salmon failed to produce meaningful results due to the limitations of the existing annotation(57). To address this, we constructed a more comprehensive transcript annotation by integrating gene models from GENCODE and NCBI’s macaque annotations (see the Comprehensive Macaque Annotation section in Methods), allowing for a broader representation of isoform diversity. Using this improved annotation, we realigned the RNA-seq reads to the genome (see Alignment section in Methods) and focused our analysis on two of the strongest predictive genes in humans, *PALM* and *SNAP25*, to assess whether they exhibit similar isoswitching trajectories in macaques.

We partitioned the macaque samples into developmental stages following the same approach used for humans. Prenatal samples were labeled as MS0, while postnatal samples were categorized into MS1 (ages 0–4), MS2 (ages 4–8), MS3 (ages 8–20), and MS4 (ages 20+) based on currently proposed developmental milestones(58).

As shown in **Figure 5e**, the same exon identified in *PALM* in humans-exon 8-exhibits minimal coverage during the prenatal stage in macaques, gradually emerging and increasing in expression over time. This exon is identical in length and results in the same protein-level amino acid changes in both species. A parallel pattern is observed for *SNAP25*, where the same exon shift leads to a nine–amino acid difference in the translated protein, consistent with our observations in humans (**Figure 5f**). These findings align with previous work by Bark *et al*.(10, 13), who documented the same *SNAP25* isoform variants in more distantly related species, including mice and chickens.

These coverage patterns provide strong evidence that the isoswitches identified in the human brain are also present in macaques. However, a larger dataset and a more comprehensive transcript annotation will be necessary for a genome-wide analysis of isoform dynamics during macaque forebrain development.

### Protein-level consequences of isoswitches with high model importance

To explore potential functional consequences of transcript-level regulation, we analyzed protein products of age-associated isoform pairs to identify examples of protein domains present at switch sites. We examined 1,000 isoform pairs whose IF trajectories were among the most predictive features for brain age. These pairs represent 1,000 high-importance genes that exhibit dominant isoswitches across the age spectrum. This expanded gene set was obtained by increasing the threshold in our feature ablation procedure from 100 to 1,000 genes (see Feature Selection section in Methods). In each iteration, we first selected the most predictive isoform, as defined in Methods. We then identified a second isoform from the same gene whose isoform fraction trajectory had the strongest negative Spearman correlation with the first, reflecting the typical opposing patterns seen in isoswitching.

For 968 protein-coding genes in the list of 1,000 high-importance genes, we compared the protein sequences encoded by each isoform pair. We confirmed that 787 / 968 isoform pairs showed divergent protein sequences, ranging from single amino acid differences to large gaps spanning more than 100 amino acids (see Table 1). Among the various ways isoforms can diverge in their protein products, we focused on pairs exhibiting a single gap, which represents the inclusion or exclusion of a continuous sequence of coding exons, likely indicating a change in a specific protein domain. These changes do not directly establish mechanistic links to aging-related phenotypes, but highlight potential protein-level consequences of the isoswitches.

**Table 1.** Difference in protein sequence lengths in proteins encoded by pairs of switching isoforms.

| Length difference (aa) | Number of pairs | Proportion (%) |
| --- | --- | --- |
| < 10 | 53 | 15.4 |
| 10–49 | 151 | 43.9 |
| 50–99 | 52 | 15.1 |
| $\geq 100$ | 88 | 25.6 |

We performed pairwise global (Needleman-Wunsch) alignment of the protein sequences encoded by the 787 isoform pairs identified earlier(59). We identified 344 pairs exhibiting a single gap, with the gap length corresponding to the length difference between the two proteins. For a large majority (≈84.6%) of instances, we observed gaps of at least 10 amino acids, with some more than 50 amino acids (see Table 1).

To annotate the protein domains affected by these gaps–hereinafter called the isoform-specific sequences–we used Foldseek(60) for fast structural alignments against large protein databases. Structure-based alignments improve search sensitivity, as protein domains can be conserved structurally despite low sequence-level conservation. We searched ColabFold-predicted structures of the longer isoforms against the CATH50 database(39, 61, 62) (see Methods). The alignments were filtered to include only those with at least 90% coverage of the isoform-specific sequence and an E-value of ≤ 0.01.

We identified 134 out of 344 isoform-specific sequences that contained a CATH domain. Notably, 96 of these 134 domains shared protein structure superfamilies (see Methods). Seven superfamilies were repeated at least five times, with examples including the P-loop containing nucleotide triphosphate hydrolases superfamily and the leucine-rich repeat variant (LRV) superfamily, which appeared 9 and 10 times, respectively. Given that CATH (v4.3.0) contains over 500,000 domains and 6,000 superfamilies, the recurrence of the same domain or superfamily is unlikely to occur by chance.

Next, we highlight two genes with isoform changes that affect a P-loop domain, illustrating how transcript-level shifts can have biological consequences. While LRV domains appeared frequently in our search results, they remain less well studied in the context of brain function.

ADP ribosylation factor-like GTPase 16 (*ARL16*) is a protein-coding gene that encodes a GTPase responsible for hydrolyzing GTP into GDP. In ARL16, we observed an inverse relationship between isoforms CHS.23550.1 and CHS.23550.15 in our earlier predictive model, with CHS.23550.1 decreasing and CHS.23550.15 increasing with age. The first 80 amino acid residues of CHS.23550.1 are missing from CHS.23550.15, and this missing region contains two P-loop domains, commonly found in ATP- and GTP-binding proteins(63) (Supplementary Figure 5a). The P-loop is responsible for phosphate binding and is essential to ARL16’s function as a GTPase. Structurally, P-loops appear as loops connecting a beta sheet to an alpha helix, and are often followed by another beta sheet (e.g., *β* – [P-loop] – *α* – *β*)(64, 65). Although the P-loop exhibits low sequence conservation, its structure is highly conserved(66), enabling reliable annotation of the structure using Foldseek.

The gene *RRAGB* (Ras-related GTP-binding B) is another example in which the dominant isoform undergoes changes in the P-loop domain. We confirmed an isoform shift from CHS.58120.2 to CHS.58120.1 for *RRAGB* and identified the 28aa residues missing from CHS.58120.2 as a P-loop domain (Supplementary Figure 5b). Like *ARL16*, *RRAGB* also encodes a GTPase, implying that changes in this domain may also affect GTP binding activity.

Previously, the shorter isoform of *RRAGB* was considered the dominant form expressed in most tissues, with the longer isoform expressed primarily in the brain(67). While the expression of both short and long isoforms of *RRAGB* has been reported in the brain(68), the interaction between their IF levels has not been explored. Here we provide a novel finding that the longer isoform (CHS.58120.1) appears to be the dominant variant in the postnatal brain.

*RRAGB* is part of the Ragulator-Rag complex, which binds to and recruits mTORC1 to the lysosome, where cellular materials get broken down into nutrients(69). mTORC1 promotes cellular anabolism, affecting processes like synaptic transmission and myelination(70, 71). *RRAGB* regulates mTORC1 in response to nutrient availability, ensuring mTORC1 persistence during nutrient deprivation to maintain vital organ function during starvation. Figlia et al. showed that each of *RRAGB*’s isoforms confers mTORC1 persistence through distinct mechanisms(68). The functional difference between these two isoforms is attributed to the 28-amino acid addition in the longer isoform, which appears to impede GTP binding. Together with our DTU results, this suggests that the mechanism of action for RRAGB changes between fetal development and the postnatal brain.

## Discussion

In this study, we demonstrate the extensive role of isoswitching in human brain development and aging. Isoform dynamics represent a layer of analysis that is often underexplored in the field in comparison to total gene expression. While previous studies have reported isolated instances of age-associated isoform variation in individual genes, we show here that this phenomenon is widespread, strongly predictive of age, and has not been systematically characterized at this scale. Many molecular and cellular processes change throughout development and aging across the brain and the broader organism; our results highlight isoswitching as one such coordinated pattern, without implying that it acts as a primary driver of these changes. We anticipate that the genes identified in this study will serve as valuable candidates for future work aimed at uncovering the underlying mechanisms.

A limitation of this study is that the relationship between these isoswitching events and the cellular composition of the brain samples remains unresolved. The consistency of the observed patterns across samples suggests that, even if these signals are influenced by changes in cell-type composition, such changes are themselves systematic and biologically meaningful. Disentangling these effects will require temporal analyses at single-cell resolution which represents an important direction for future work.

Another limitation is the cross-sectional nature of postmortem brain data. Because only a single sample is available per individual, rather than repeated measurements over time, the ability to model within-individual variability is limited.

Finally, it will be important to evaluate whether these isoform dynamics are specific to the brain or are observed more broadly across tissues, as well as to determine whether similar patterns are present across different brain regions.

## Methods

### Pairwise global alignment of protein sequences

We performed pairwise global alignment of two protein sequences using the pairwise-sequence-alignment Python module at https://github.com/aziele/pairwise-sequence-alignment. The module implements the Needleman-Wunsch algorithm for global sequence alignment and is part of the EMBOSS package(59).

For each alignment result, we checked whether the longest gap matched the length difference between the two sequences, excluding cases where the original sequences were of identical length. For all such cases, we saved the isoform-specific sequence and noted which of the two sequences was longer.

### Structure-based domain searches

To search for protein domains intersecting our list of isoform-specific sequences, we used Foldseek(72) to align the predicted protein structures of the longer isoforms against the CATH50 database, which we obtained using the foldseek databases command. CATH50 comprises representative domain structure from CATH(62) (v4.3) and AlphaFold DB(73), clustered at 50% sequence identity. Most of the structures were downloaded from <u>isoform.io</u>, a public online database hosting ColabFold-predicted protein structures for 237,275 human isoforms(74). We folded 36 isoforms that were omitted from the database due to their length (>2,000aa), using ColabFold on a Nvidia A100 GPU(39, 61). CHS.53345.5 was the only isoform that could not be folded due to its excessive length (>4,000aa).

We filtered the Foldseek results based on their E-values and alignment coordinates, requiring an E-value no larger than 0.01 and a minimum query coverage of 90%. Query coverage was calculated as the proportion of the isoform-specific sequence length that aligned. These filters ensured that only statistically significant domain hits with meaningful overlap in the gap region were retained.

We compiled two versions of the filtered Foldseek results: one including all domain hits and another containing only the top hit for each query. The latter was used to assess the enrichment of specific CATH superfamilies. The filtered output includes alignment coordinates, CATH domain ID, CATH superfamily ID, query coverage, E-value, and bit score for each hit. When multiple hits were identified for a single query, the domain with the lowest E-value was selected, with ties resolved arbitrarily. Domain and superfamily annotations were primarily obtained using the Foldseek-provided cath-v4_3_0.alphafold-v2.2022-11-22.tsv file. In cases where this information was unavailable, we referred to the cath-domain-list.csv file from the official CATH classification data available on the CATH website(62).

### Stacked model

Let *T* = {*s*_1_, *s*_2_, … *s*_*N*/2_} be the set of all training samples where *N* is the total number of brain samples. Each sample *s*_*i*_ has an isoform fraction (IF) vector of *X*_*i*_ = [*f*_*t*_1__ *f*_*t*_2__ … *f*_*t_p_*_] with *p* elements where *f*_*t_j_*_ represents the IF value from a selected transcript *t*_*j*_. After feature selection, in each vector *X*_*i*_ there remained a total number of *p* = 341 transcripts from the 100 selected genes. For a training sample *s*_*i*_, the random forest regressor predicts an age based on its *X*_*i*_, which we will call *s*_*i*_*′*, and this *s^′^* will fall within the boundaries of either 1 or 2 of the overlapping lasso regressor windows:

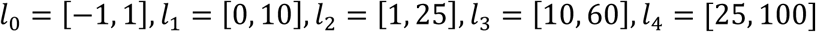

Each of these lasso models is trained exclusively on the subset of samples corresponding to its respective age range, as determined by the random forest predictions. For example, *l*_0_ will be trained on 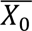 where 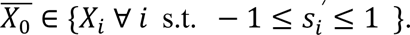 The random forest regressor was constrained to have a minimum of 10 samples per leaf prior to the second regression step. This parameter was retained in all subsequent steps, as it proved effective in improving model generalization. It is important to note that all steps in both regressor layers use log-ages as input and output. However, corresponding real ages are used here for clearer presentation of the model structure.

We selected lasso regression for this second layer due to its inherent ability to perform feature selection while strictly penalizing overfitting. Unlike standard regression models, lasso applies L1 regularization, which forces some regression coefficients to shrink to zero, effectively removing less relevant features from the model:

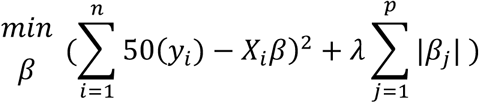

where *y*_*i*_ is the log-age of the subject in sample *s*_*i*_, *X*_*i*_ is the IF vector of sample *s*_*i*_, *p* is the number of features, *β* is the vector of regression coefficients, and *λ* is the regularization parameter, which we set to 0.1. Log-ages were multiplied by an arbitrary factor of 50 to amplify age differences for clearer separation across regressors.

### Feature selection

Feature selection presented several challenges because we wanted to avoid discarding features (genes) that might be important. We considered using backward elimination, where features are removed based on the decrease in model accuracy when a given feature is excluded from the training set(78). If two features exhibit a highly similar association with the target variable, eliminating one has little impact on performance. Likewise, in forward selection, features are added based on their unique contribution to predictive power, meaning that strongly correlated features provide minimal additional information(78, 79).

However, removing redundant isoform fractions solely based on redundancy is not ideal for our analysis. Consider a scenario where Gene A undergoes a linear isoform shift with aging, where isoform A₁ increases while A₂ decreases. If an entirely independent Gene B exhibits a similar pattern, with isoform B₁ increasing and B₂ decreasing, conventional feature elimination methods would likely discard one of these genes, as their predictive information is considered redundant.

Yet, if both genes undergo equally significant isoform shifts, capturing both is crucial for a comprehensive understanding of developmental and aging processes. Furthermore, whether these genes are functionally related within the same pathway or entirely independent, there is no way to determine a priori which isoform shift is more biologically relevant. Prematurely discarding one would risk losing valuable biological insight, emphasizing the need for a feature selection approach that preserves meaningful signals for downstream analysis.

Given these considerations, we chose to identify the strongest predictive genes one by one, as outlined in the Results section. The feature ablation approach(44), used to examine how predictive power is distributed across genes, also proved to be an effective method for feature selection. Starting with the most important features, we iteratively removed all isoforms associated with the top-ranked gene and retrained the model at each step. Although the procedure was intended to continue until model performance declined to a suboptimal level, this threshold was not breached even after the removal of a substantial number of genes (**Figure 4a**). We ultimately selected the top 100 predictive genes for inclusion in our decision forest model-an arbitrary cutoff chosen to constrain the scope of downstream biological interpretation. The top 5000 genes ordered based on predictive value are provided in Supplementary Table 1.

All training and testing splits were stratified to ensure a balanced distribution of samples across age ranges. To achieve this, we defined custom age bins based on the distribution and constraints of our dataset. These bins were more finely spaced in earlier developmental stages and were defined as follows: from −1 to −0.3 in intervals of 0.1, from −0.3 to 0 in a single 0.3 interval, from 0 to 1 year in intervals of 0.2, from 1 to 3 years in 1-year intervals, from 3 to 30 years in 3-year intervals, and from 30 to 100 years in 10-year intervals. This stratification step was kept in all subsequent models.

At each iteration, we trained a model on the LIBD training set using log-ages and identified the most important feature based on Gini scores. The feature selection procedure was not informed by the LIBD test set or the independent test data. Each random forest regressor consisted of 10 estimator trees, with a minimum of 10 samples required per leaf. The top feature identified at each step corresponded to an isoform fraction vector associated with a parent gene. We then removed all isoforms associated with that gene and repeated the process. Additionally, to assess the influence of log transformation, we trained separate models using both raw and log-age values at each step, comparing their R² scores to evaluate potential differences in predictive power. We observed minimal impact on the model’s predictive accuracy, and no significant differences between the R² trends of the log-scaled and raw age models (**Figure 3a**).

In order to assess the stability of the top selected genes, we performed feature selection across 10 random splits of the dataset. For each split, we identified the top 1000 features and compared their overlap across splits. Between all pairwise combinations of splits, we calculated the Jaccard distance. The mean Jaccard distance for the top 100 genes was 0.43 (min: 0.31, max: 0.53), and for the top 1000 genes it was 0.64 (min: 0.59, max: 0.67). The top 513 most frequent genes are present across all splits, which we provide in Supplementary Table 2. Model performance through the feature ablation process appears consistent across splits (Supplementary Figure 4).

### Isoform quantification and filtering

RNA-seq reads were quantified at the isoform level using Salmon(80) (version 1.10.0) with the CHESS 3(24) (v.3.1.1) annotation, and scaled using *tximport*(81). Transcripts were then filtered using the SPIT(25) pre-filtering module to retain reliably expressed, multi-isoform genes. For DTU analysis, we defined the prenatal group as the control and the postnatal group as the case, allowing us to capture potential subgroup-specific isoform shifts across the wide postnatal age range. To further examine splice site usage and transcript structure, we performed splice-aware alignments using HISAT2(82) and generated aggregated coverage profiles for visualization. RNA quality was consistently high across samples and was assessed using standard QC procedures. For macaque samples, we used the Mmul_10(83) assembly together with a more comprehensive merged annotation to improve isoform resolution. Full details on quantification, filtering criteria, alignment, and annotation are provided in the Supplementary Methods.

## Data availability

The source code supporting all modeling and statistical analyses, along with processed data, selected gene sets, and structural domain search results, has been deposited in Zenodo: https://doi.org/10.5281/zenodo.15335237

## Acknowledgements

We would like to acknowledge Martin Steinegger and Milot Mirdita for their contributions to the functional analysis pipeline. We also thank Ales Varabyou for assembling the *Macaca mulatta* transcriptome annotation.

This work was supported in part by the U.S. National Institutes of Health under grants R01-MH123567, R01-HG006677, and R35-GM156470.

