## Supplementary Figures for "Isoswitching in human brain development and aging"

**a.**

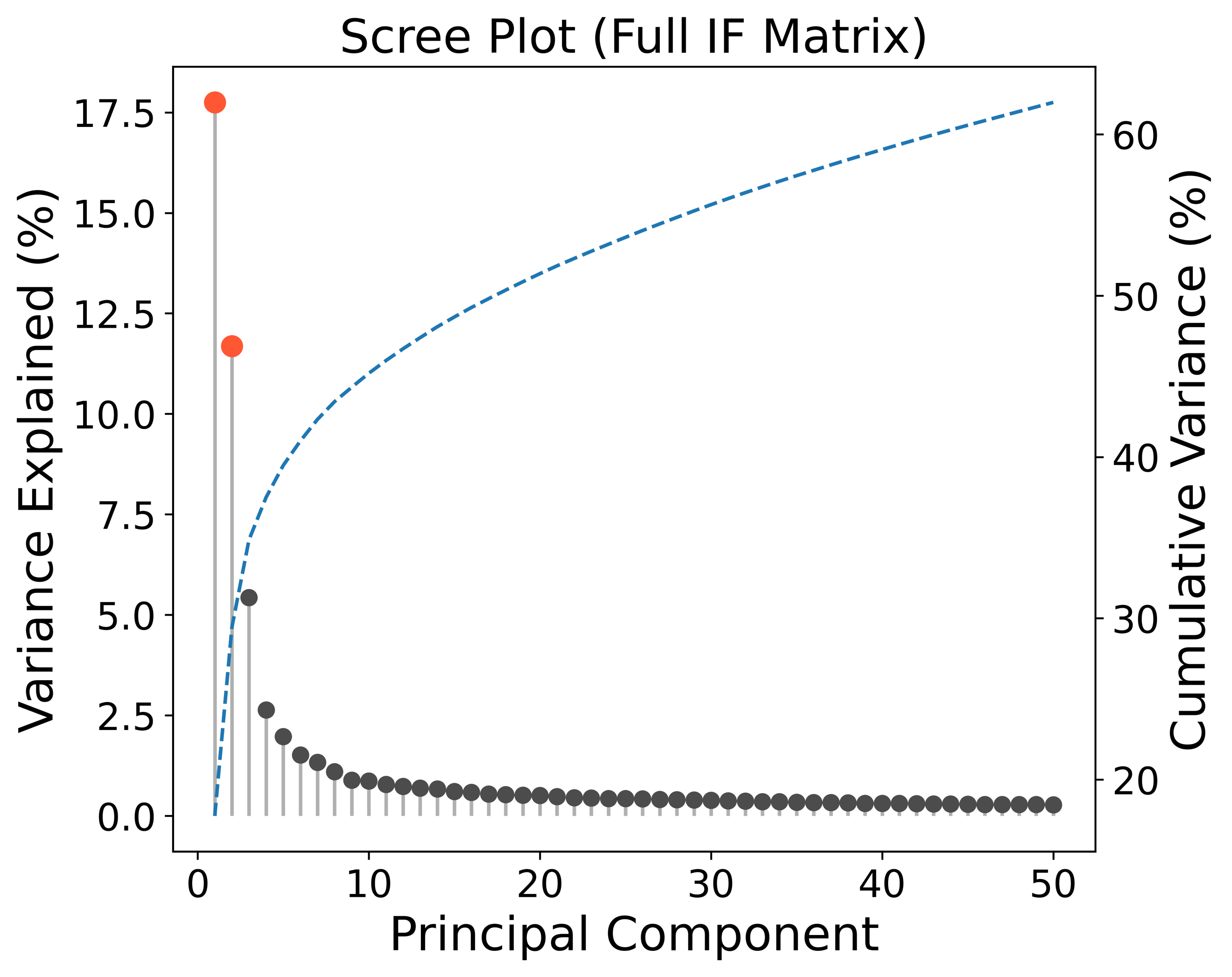

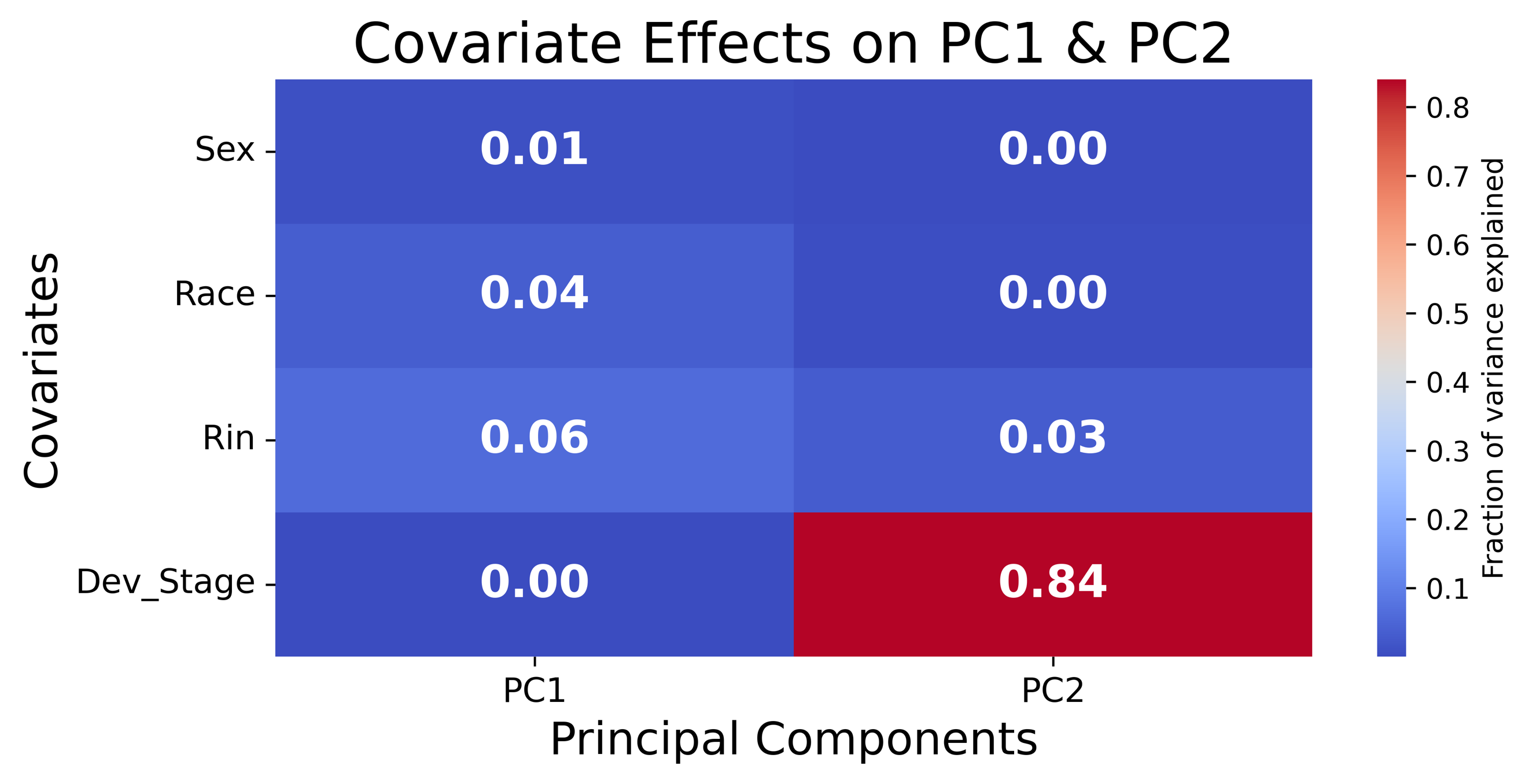

**Supplementary Figure 1.**

**a.** Scree plot from the PC analysis of all samples (n=341) and isoforms (n=29,757) shown in Fig. 1b. Variance explained by the top 50 principal components is shown, along with cumulative variance. PC1 explains 17.75% of the variance and PC2 explains 11.69%, with a cumulative variance of 29.44% for the first two components. **b.** Covariate effects on PC1 and PC2 are measured using Type II ANOVA on linear models with each PC as the response variable. Developmental stage (Dev_Stage) represents pre- vs. postnatal stage. Categorical variables (sex, race, and developmental stage) were modeled as factors.

**b.**

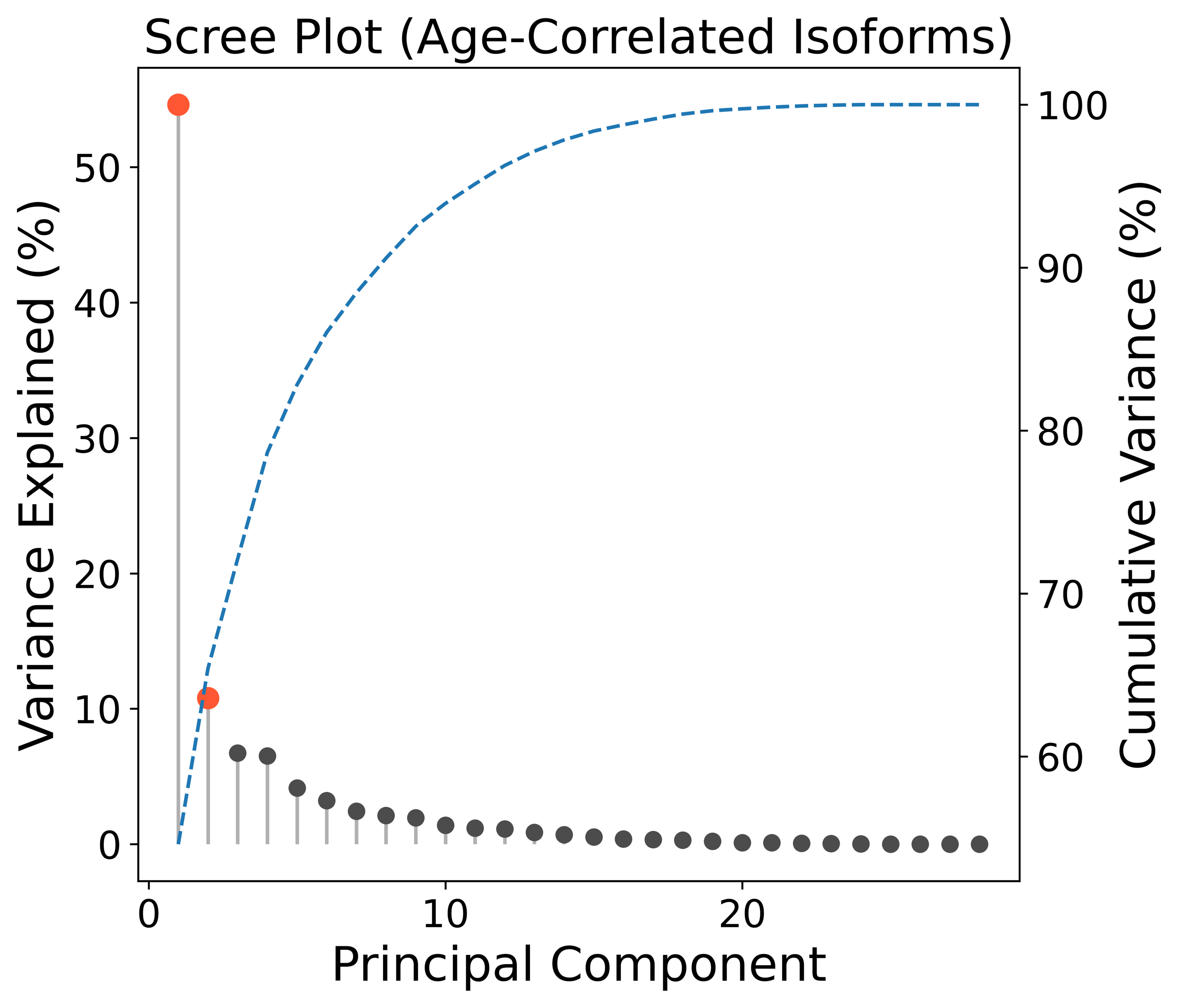

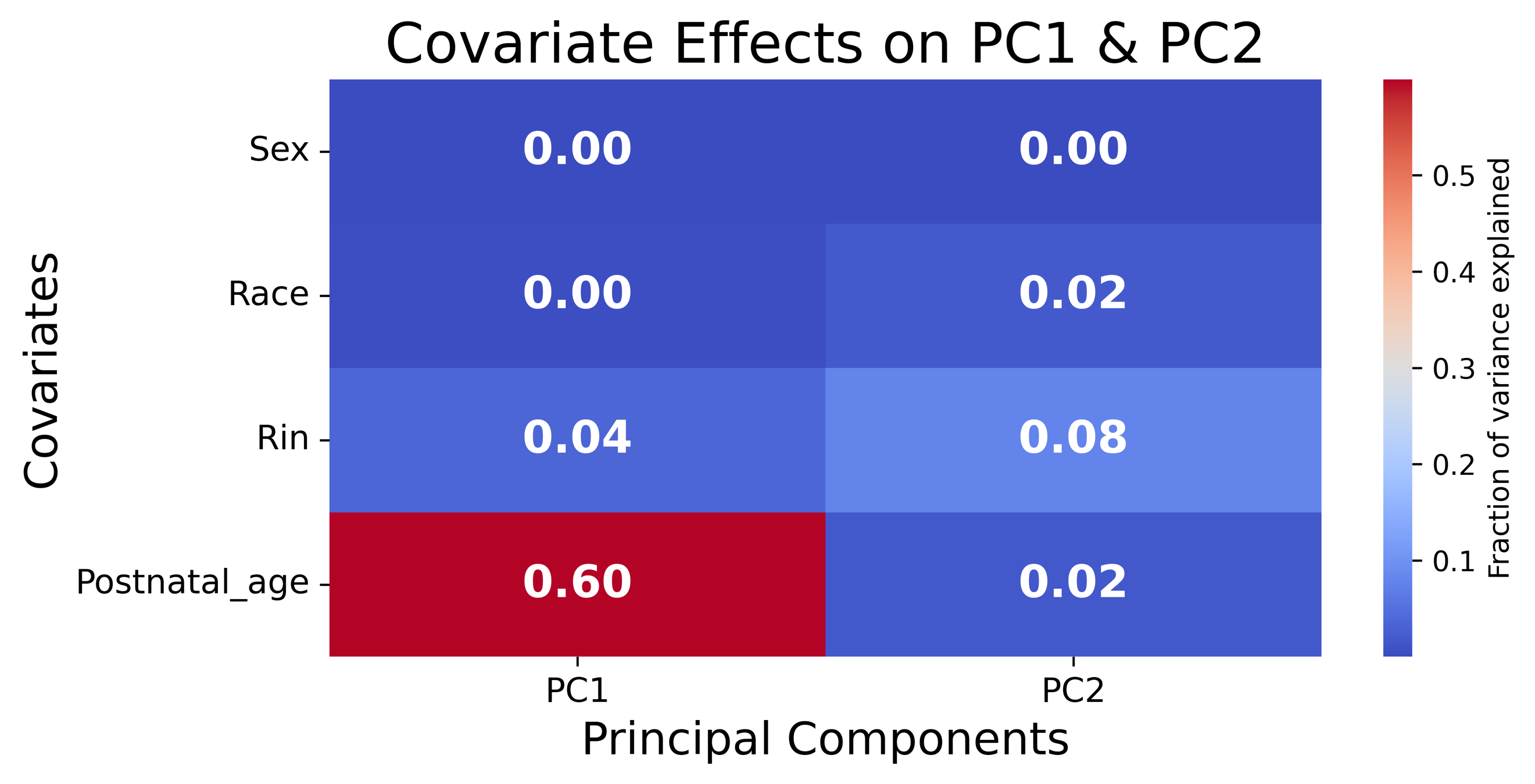

**Supplementary Figure 2.**

**a.** Scree plot from the PC analysis of postnatal samples (n=285) and the top age-correlated isoforms (n=28) shown in Fig. 1f. Variance explained by all 28 principal components is shown, along with cumulative variance. PC1 explains 54.61% of the variance and PC2 explains 10.79%, with a cumulative variance of 65.40% for the first two components. **b.** Covariate effects on PC1 and PC2 are measured as reported in Supplementary Figure 1b.

**a.**

**b.**

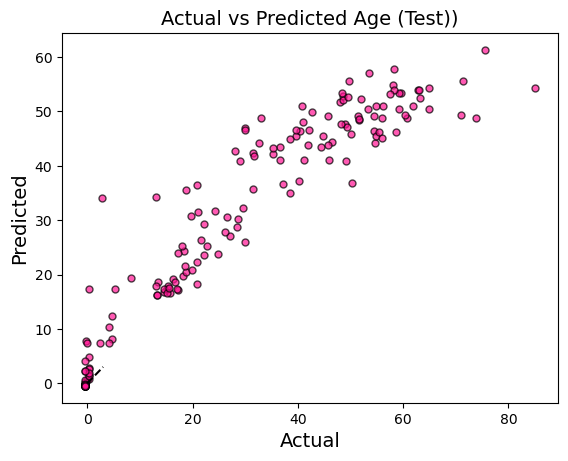

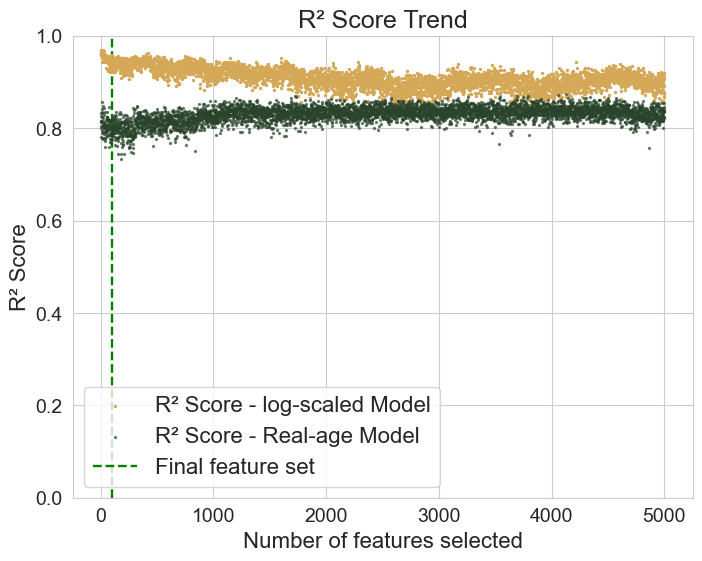

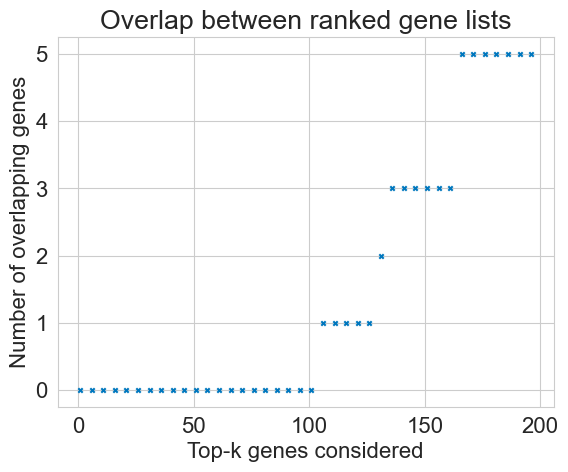

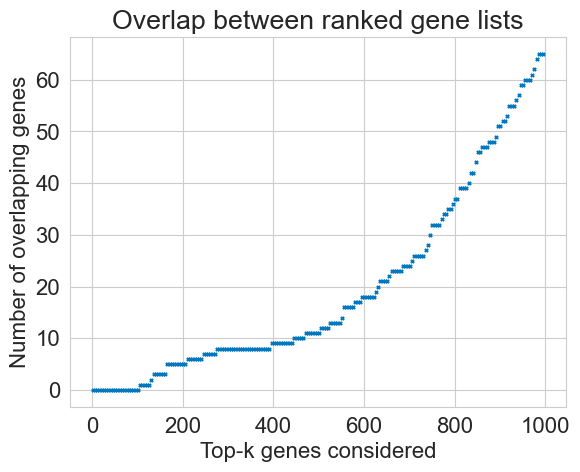

**Supplementary Figure 3.**

**a.** Actual vs. predicted ages from the initial total gene expression random forest regressor trained on 18,637 genes (Training: R² = 0.99, RMSE = 12.25; Testing: R² = 0.88, RMSE = 7.86). **b.** R² scores throughout the feature ablation process. At each step, the most important gene from the previous model was removed (without replacement), and the model was retrained. The model trained on log-transformed ages is shown in yellow, and the model trained on raw age values is shown in green. **c, d.** Overlap between top predictive genes from IF and gene expression models as a function of k, the number of top-ranked genes considered.

**a.**

**b.**

**c.**

**d.**

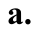

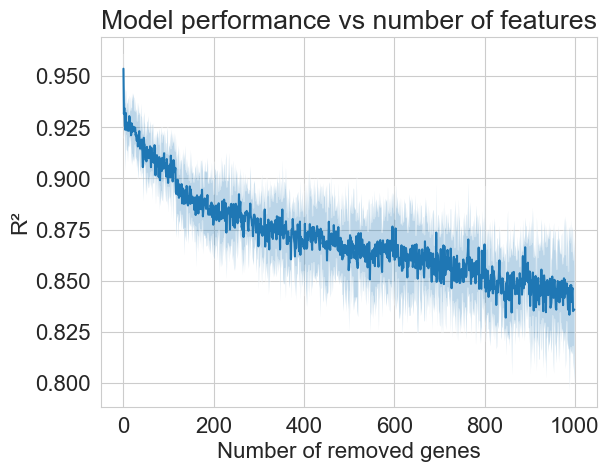

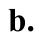

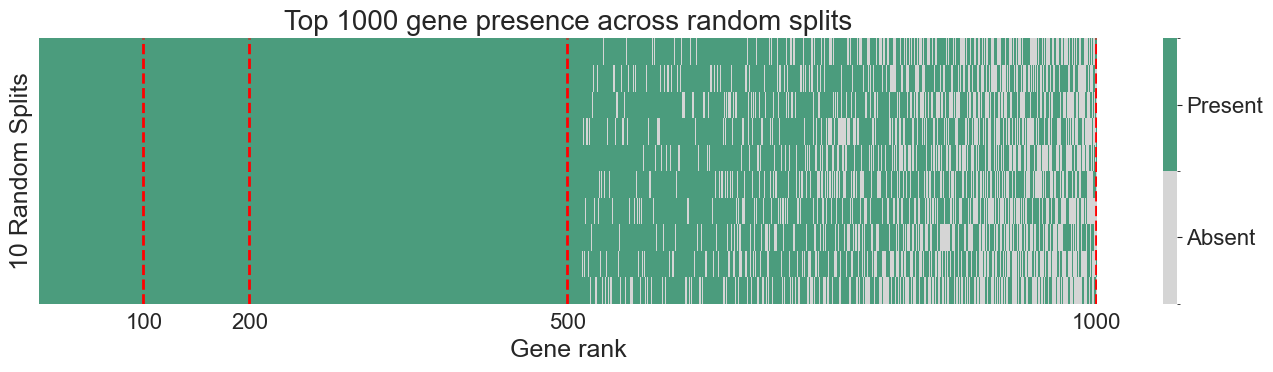

**Supplementary Figure 4.**

**a.** R² scores across the feature ablation process over 10 random splits of the data. For each number of genes removed, the shaded region represents ±1 standard deviation across splits, while the dark blue line indicates the mean trend. **b.** Heatmap showing the presence of top-ranked genes across splits. Genes are ordered by frequency of occurrence, with the top 514 genes appearing in all 10 splits.

**Supplementary Figure 4.**

**a.** Actual vs. predicted ages from the initial total gene expression random forest regressor trained on 18,637 genes (Training: R² = 0.99, RMSE = 12.25; Testing: R² = 0.88, RMSE = 7.86). **b.** R² scores throughout the feature ablation process. At each step, the most important gene from the previous model was removed (without replacement), and the model was retrained. The model trained on log-transformed ages is shown in yellow, and the model trained on raw age values is shown in green. **c, d.** Overlap between top predictive genes from IF and gene expression models as a function of k, the number of top-ranked genes considered.

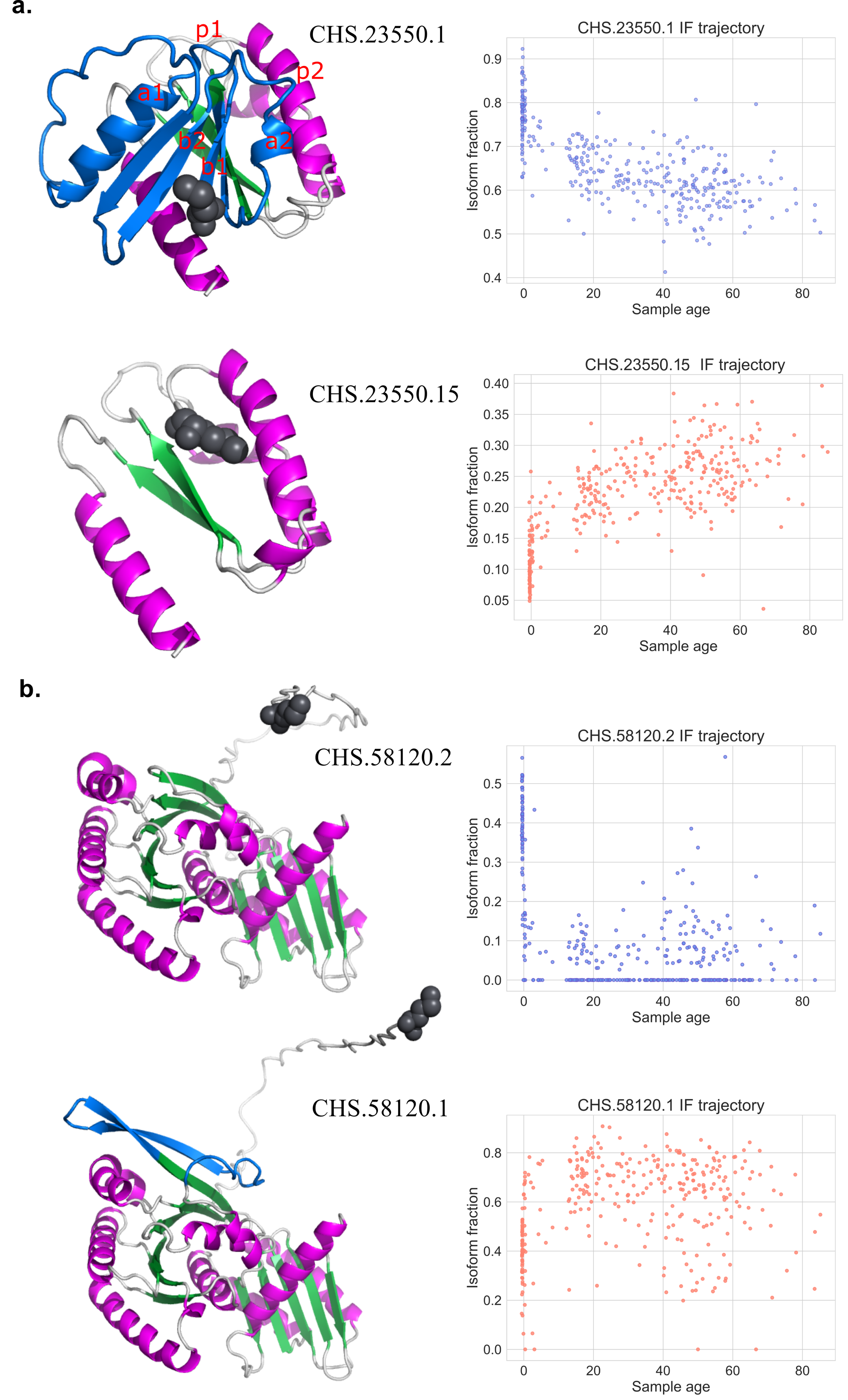

**Supplementary Figure 5.**

Aligned protein structures and isoform trajectories for ARL16 and RRAGB. Structural regions are colored: loop (light grey), alpha helix (magenta), beta sheet (green), and isoform-specific regions (blue). N-termini are marked by dark grey spheres. **a.** Left: ARL16 isoform CHS.23550.1 contains two additional P-loop domains (blue) compared to CHS.23550.15. P-loops are labeled p1 and p2 in red. Right: Corresponding IF trajectories for CHS.23550.1 (blue) and CHS.23550.15 (orange). **b.** Left: RRAGB isoform CHS.58120.1 (right) shows an elongation and added beta sheet (blue) relative to CHS.58120.2. Right: IF trajectories for CHS.58120.2 (blue) and CHS.58120.1 (orange).
