## Supplementary Methods for "Isoswitching in human brain development and aging"

**Gene expression model**

Transcript counts were summed across isoforms for each gene to obtain a gene-level count matrix for the LIBD data. This matrix was normalized to counts per million (CPM) and filtered to retain genes with expression of at least 1 CPM in at least 34 samples (10% of the dataset) which resulted in a 18,637 genes. Consistent with differential gene expression analysis protocols(75), we further normalized this count matrix using the trimmed mean of M-values (TMM) method(76, 77).

We used the filtered and normalized CPM matrix as input to train a random forest regressor to predict log-scaled ages of the LIBD samples. We then performed iterative feature ablation, removing the top predictive gene at each step, to select the top 5000 predictive genes (Supplementary Figure 3).

**Isoform quantification and filtering**

RNA-Seq reads from all brain samples were mapped to transcripts in the CHESS 3(24) (v.3.1.1) gene catalog of the GRCh38 human genome (p.12) using Salmon(80) (version 1.10.0). For Salmon, the entire genome was used as a decoy sequence, and quantification was performed with the following options:

--validateMappings --mimicBT2 --recoverOrphans --gcBias.

Isoform quantifications were extracted as Transcripts Per Million (TPM) values and scaled using the "dtuScaledTPM" with the R package *tximport(81)*. These scaled read counts were used as the unit of expression in all downstream analyses. Quantified transcripts were filtered prior to DTU analysis with the SPIT pre-filtering module which applies the following criteria in order:

- An isoform must be expressed at a level of at least 1 CPM in a minimum of $n_{small}$samples, where $n_{small}$ is set to the default 12.
- An isoform must have a positive read count in at least 20% of samples in both groups of analysis.
- A gene must have a minimum read count of 10 in at least 10 samples.
- An isoform must have an IF value greater than 0.1 in at least $n_{small}$samples.
- Following these filtering steps, a gene must be left with more than 1 isoforms.
- For each gene, the control group should exhibit a dominant isoform. This is measured by the same isoform having the highest IF in at least 50% of control samples.

In our analysis of transcriptomic changes across brain development and aging, we designated the prenatal group as the control and the postnatal group as the case. This setup was particularly suitable for SPIT, which is designed to identify potential substructures within the case group. Given the wide age range in the postnatal cohort -from infancy to late adulthood- isoform shifts may be more pronounced within specific age subsets than across the entire group. SPIT is capable of detecting such subgroup-specific patterns by accounting for bimodal or multimodal distributions within the case group, thereby preserving signals that might otherwise be overlooked in traditional DTU analyses. While changes in isoform usage are also expected during prenatal development, our dataset is largely composed of samples between 15 and 20 post-conception weeks, limiting the ability to detect meaningful clustering or temporal substructure within the prenatal group.

For the GTEx heart samples, RNA-Seq reads were again quantified using Salmon(80) with the RefSeq annotation (release 110). As this analysis did not involve a case–control design, we focused on including only the most stably expressed genes to minimize noise in the predictive model. Genes were filtered based on their median CPM across all samples, retaining the top 1,000 genes with at least two annotated isoforms. Varying this threshold to 500 or 5,000 genes did not result in appreciable changes in model performance.

**Alignment**

To assess transcript coverage and examine stage-specific inclusion or exclusion of splice sites, we performed read alignments using HISAT2(82) in “splice-aware” mode for both human and macaque datasets. Human samples were aligned to the GRCh38.p12 reference genome, while macaque samples were aligned to the Mmul_10(83) genome assembly.

To generate a composite view of consistent alignment patterns across multiple samples, we used TieBrush(84) to aggregate alignments, followed by TieCov to compute coverage profiles. The resulting coverage data were then filtered, normalized, and visualized using the Integrative Genomics Viewer (IGV)(85) to provide a comprehensive picture of isoform usage across developmental stages.

**RNA-Sequencing and quality control**

Tissue collection and sequencing protocols for the brain samples from LIBD have been previously described by Lipska *et al*(86). and Jaffe *et al*(87). The median RNA integrity number (RIN) of the control samples used in our analysis was 8.5, with over 80% of the samples having RIN values of 8 or higher. Samples with RIN values below 5 were excluded from the analysis. Additionally, we assessed the quality of the RNA-seq reads using FastQC(88) and MultiQC(89), which showed consistently high results, with 98% of our samples having the majority of their reads with a median per-base sequence quality score above 30.

**Comprehensive macaque annotation**

To process rhesus macaque samples, we used the Mmul_10 genome assembly(83, 90-92), which was the most contiguous and accurate assembly available at the time of the study. Although several alternative gene catalogues for Mmul_10 are available, substantial differences in covered loci and alternative splicing events made those inadequate for our analysis. Instead, we chose to adopt a more comprehensive genome annotation that will be released in a forthcoming study. This annotation represents a union of the Ensembl and NCBI annotations for the Mmul_10 assembly. This combined annotation comprises 158,009 transcripts and 50,083 genes, encompassing both protein-coding and non-coding biotypes.
